# Assessing the *in planta* efficacy of oxathiapiprolin as a potential treatment for kauri dieback disease

**DOI:** 10.1101/2024.12.01.626252

**Authors:** Hannah F. Robinson, Jade T. T. Palmer, Monica L. Gerth

**Affiliations:** School of Biological Sciences, Victoria University of Wellington, Wellington, New Zealand

**Keywords:** oxathiapiprolin, *Agathis australis*, kauri dieback, *Phytophthora agathidicida*, plant trial, Zorvec

## Abstract

*Phytophthora agathidicida*, a plant pathogenic oomycete, causes fatal dieback disease in New Zealand kauri trees (*Agathis australis*). Currently, few treatments exist to prevent or cure this infection. Previous research has demonstrated the potent *in vitro* inhibition of multiple lifecycle stages of *P. agathidicida* by the oomycide oxathiapiprolin. In this study, we have evaluated the efficacy of oxathiapiprolin *in planta* as either a protective or curative treatment. Kauri seedlings (1–2 years old) were treated with 10 or 50 mg of oxathiapiprolin, in the form of Zorvec Enicade, *per* seedling as a soil drench either before (7 days) or after (15 days) inoculation with *P. agathidicida* NZFS 3770 to test for protective and curative activities, respectively. Results showed that oxathiapiprolin treatments successfully protected the kauri seedlings from disease, with the higher dose (50 mg) demonstrating greater efficacy. However, the treatments did not cure kauri seedlings already infected with*P. agathidicida*, likely because the infection was already well-established by the time of treatment. This study demonstrates that, while oxathiapiprolin shows protective effects against *P. agathidicida* infection in kauri seedlings, its lack of curative properties significantly limits its potential as a practical tool for managing kauri dieback disease.

## Introduction

*Phytophthora* species are plant pathogenic oomycetes that destroy crops and natural ecosystems worldwide. Their complex, multi-spore lifecycle and the absence of common targets of antifungal treatments make them challenging to control chemically (Judelson and Blanco, 2005; Giachero, Declerck and Marquez, 2022). Oxathiapiprolin, marketed as DuPont™ Zorvec™, is the first member of a novel class of oomycides (piperidinyl thiazole isoxazolines) and exhibits high activity against *Phytophthora* species, providing both protective and curative effects (Ji *et al*., 2014; Matheron and Porchas, 2014, 2015; Ji and Csinos, 2015; Miao *et al*., 2016; Pasteris *et al*., 2016; Qu *et al*., 2016; Bittner and Mila, 2017; Cohen, Rubin and Galperin, 2018, 2019; Hao *et al*., 2019; Cohen, 2020; Cohen and Rubin, 2020). Therefore, it has potential as a treatment for currently uncontrolled or difficult-to-manage *Phytophthora* diseases.

In New Zealand (NZ), *Phytophthora agathidicida* is the causative agent of dieback disease in endemic kauri (*Agathis australis*) (Bradshaw *et al*., 2020). Kauri are large, long-lived conifer trees of ecological and cultural significance (Mainwaring, Vink and Gerth, 2023). Kauri dieback is a lethal root rot disease characterised by the hyperproduction of resin around the tree collar and lower trunk, leaf yellowing, and canopy thinning, typically resulting in death within 10 years (Bellgard *et al*., 2016; Bradshaw *et al*., 2020). The now widespread nature of dieback disease within the natural range of kauri in the northern North Island threatens the health of kauri forests, making control of the epidemic imperative (Bradshaw *et al*., 2020).

Current strategies for managing kauri dieback include containment measures and chemical controls (Bradshaw *et al*., 2020). Containment efforts involve installing footwear washing stations at track entrances, upgrading or closing tracks in vulnerable areas, and implementing restrictions on earthworks, stock grazing, and animal release in kauri lands (Bellgard *et al*., 2016; Bradshaw *et al*., 2020; Kiro, 2023). Chemical control trials have shown that phosphite injections into the trunks of infected kauri trees have a curative effect (Horner and Hough, 2013; Horner, Hough and Horner, 2015; Horner and Arnet, 2020).

However, phytotoxicity at higher phosphite concentrations and the potential for resistance warrant research into alternative chemical controls with greater potency and tolerability (Dobrowolski *et al*., 2008; Horner, Hough and Horner, 2015; Horner and Arnet, 2020).

Oxathiapiprolin is a promising candidate for the chemical control of *P. agathidicida*. It can be applied as a soil drench to manage soilborne diseases, such as kauri dieback, and its acropetal and xylem systemic movement protects growing leaves and new foliage from infection (Pasteris *et al*., 2019). Oxathiapiprolin targets the oomycete oxysterol binding protein, which plays roles in sterol transport, lipid metabolism, and lipid membrane composition, impacting signalling and vesicle transport (Weber-Boyvat *et al*., 2013; Pasteris *et al*., 2016, 2019). Thus, oxathiapiprolin inhibits multiple stages of the pathogen’s lifecycle and has specific anti-oomycete activity, minimising off-target effects. Specificity is especially important for chemical controls in kauri habitats, and studies have confirmed that oxathiapiprolin has favourable environmental and toxicological profiles (Pasteris *et al*., 2016, 2019).

In previous work, our lab group has reported the potent *in vitro* inhibition of the three major lifecycle stages of *P. agathidicida* (mycelial growth, zoospore germination, and oospore germination) by oxathiapiprolin at sub-microgram *per* millilitre concentrations (Lacey *et al*., 2021). The inhibition of zoospore and oospore germination indicates that oxathiapiprolin may prevent the infection of kauri by *P. agathidicida*, while the inhibition of mycelial growth indicates curative properties. Therefore, in this study, we conducted plant trials with kauri seedlings to determine the optimal dosage of oxathiapiprolin and assess its protective and/or curative effects following inoculation with *P. agathidicida*. This paper summarises the *in planta* activity of oxathiapiprolin when applied as a soil drench.

## Materials and Methods

### P. agathidicida Isolate and Culture Conditions

*P. agathidicida* NZFS 3770 (ICMP 17027 holotype) was provided from Scion’s culture collection (Rotorua, NZ) but was originally isolated in NZ from Great Barrier Island (Guo *et al*., 2020). Upon receipt, it was initially cultured on oomycete-selective PARP agar plates composed of cornmeal agar (17 g/L; BD BBL™, Sparks, MD, USA) supplemented with pimaricin (0.001% w/v; Sigma-Aldrich, St. Louis, MO, USA), ampicillin (250 µg/mL; GoldBio, St. Louis, MO, USA), rifampicin (10 µg/mL; Sigma-Aldrich), and pentachloronitrobenzene (100 µg/mL; Sigma-Aldrich). Subsequent cultures were passaged by transferring an agar plug with a cork borer from the leading edge of mycelial growth onto 20% v/v aq clarified V8 (cV8) agar plates. Clarified V8 agar (1 L) was prepared by mixing 200 mL V8 Original Vegetable Juice with 200 mL distilled, deionized water (ddH_2_O) and 2 g CaCO_3_ for 30 min and then clarifying *via* centrifugation at 7000*g* for 10 min. Bacteriological agar (15 g; New Zealand Seaweeds, Opotiki, NZ) and 600 mL ddH_2_O were added to the supernatant, which was thereafter sterilized by autoclaving. All cultures were grown in the dark at 22 °C.

### Preparation and Maintenance of Kauri Seedlings

For the protective plant trial, kauri (*Agathis australis*) seedlings (1–2 years old, 30–70 cm tall) obtained from ōtari-Wilton’s Bush (Wellington, NZ) after germination and subsequently grown in a glasshouse were re-potted in horticultural polypropylene pots (2 L/15 cm diameter) using Tui Essentials All Purpose Potting Mix. The re-potted seedlings were left in the glasshouse for ≥ 1 week to ensure they were undamaged from the re-potting. For the curative plant trial, kauri seedlings (approximately 2 years old, 50–80 cm tall, 1.5 L pots) purchased from Naturally Native (Tauranga, NZ) were left in the glasshouse for ≥ 1 week to acclimatize. Before beginning the plant trials, all seedlings were transferred to the PC2 plant laboratory, in which they were allowed to equilibrate for ≥ 1 week. The plant lab was maintained at 18–22 °C (35–75% relative humidity). The seedlings were placed in individual aluminum trays and watered every 2–3 days until damp but not waterlogged. They were grown under fluorescent tube lights (Growshop 4 ft 4 × 54 W T5 Fluorescent Tube Fitting c/w 6500k) on a 12 h on: 12 h off cycle.

### Treatment Preparation and Application

Zorvec® Enicade® (ZE), containing 100 g oxathiapiprolin/L as an oil dispersion, was gifted by Corteva™ Agriscience (New Plymouth, NZ) and stored at room temperature. Three seedling replicates were prepared for each treatment and control type. Treatment types included seedlings inoculated with *P. agathidicida* (PA) and treated with 10 or 50 mg oxathiapiprolin/seedling (+PA, +ZE), and control types included seedlings that were uninoculated and untreated (-PA, -ZE), uninoculated but treated with 50 mg oxathiapiprolin/seedling (-PA, +ZE), and inoculated but untreated (+PA, -ZE). Seedlings were treated with 10 or 50 mg oxathiapiprolin/seedling by diluting Zorvec® Enicade® in sterile ddH_2_O to final concentrations of 0.2 or 1 mg oxathiapiprolin/mL (total volume = 50 mL), respectively, and pouring each 50 mL treatment in the root zone of the corresponding seedling, as in Horner and Hough (2013). Each pot was placed in a plastic bag that was tied around the bottom of the stem to reduce evaporation, and the seedlings were not watered for 1 week to allow time for the uptake of each treatment. Sterile ddH_2_O (50 mL) was applied to the untreated control seedlings (±PA, -ZE). For the protective plant trial, the seedlings were treated 1 week prior to the first inoculation, and for the curative plant trial, the seedlings were treated 1 day after the third (and final) inoculation (*i*.*e*. 15 days after the first inoculation).

### Preparation of Inocula and Kauri Seedling Inoculation

The inocula were prepared as follows: agar plugs (4 mm^2^) from the leading edge of *P. agathidicida* NZFS 3770 mycelial growth on 20% v/v cV8 agar were incubated in 10 mL of 2% v/v aq cV8 broth at 22 °C in the dark for 2 days. For the uninoculated control seedlings (-PA, ±ZE), blank 20% v/v cV8 agar plugs (*i*.*e*. no *P. agathidicida* present) were prepared in the same manner. As in Byers *et al*. (2021), kauri seedlings were inoculated with 5 mycelial mats (or blank cV8 agar plugs for the uninoculated control seedlings) as follows: 8 mm diameter wooden dowels were used to create 5 evenly spaced, 5–10 cm deep holes in the soil around the seedling within the root zone. The 10 mL mycelial cultures (or blanks) were poured into the holes and then covered with soil, and the seedlings were watered daily until total saturation for 1 week to promote zoospore motility and stimulate infection. The seedlings were re-inoculated twice with 3 mycelial mats (or blank cV8 agar plugs) after 1 and 2 weeks to ensure infection and then saturated with water overnight. The seedlings were monitored weekly by taking pictures, measuring seedling height, and recording observations.

### Rating Disease Development

After 6 weeks post-inoculation, the seedlings were randomized, and 5 blinded, independent assessors rated seedling health for all 15 seedlings according to the criteria listed below:

*Wilting:* assessment of the overall severity of leaf wilting for each seedling

| Rating | Description |
| --- | --- |
| 0 | no wilting |
| 1 | mild wilt symptoms |
| 2 | moderate wilt symptoms |
| 3 | severe wilt symptoms |

*Color:* assessment of the overall extent of browning/yellowing in the leaves of each seedling

| Rating | Description |
| --- | --- |
| 0 | no browning/yellowing (*i.e.* all leaves green) |
| 1 | mild browning/yellowing |
| 2 | moderate browning/yellowing |
| 3 | severe browning/yellowing |

*Dryness:* estimation of the percent of leaves (or total leaf material) that have dried out in each seedling

| Rating | Description |
| --- | --- |
| 0 | 0% |
| 1 | > 0% – ≤ 10% |
| 2 | > 10% – ≤ 50% |
| 3 | > 50% – < 100% |
| 4 | 100% |

*New Foliage:* observation of the presence of new, un/healthy growth

| Rating | Description |
| --- | --- |
| 0 | yes – new, healthy growth is present |
| 1 | no – no new growth is present or new growth is unhealthy |

The results were analyzed on GraphPad Prism 8.4.3 using a 2-way factorial ANOVA with the Tukey *post hoc* correction to determine significant differences (*p* ≤ 0.05). To establish that the assessors were able to use the criteria to come to agreement on wilting (scored 0–3), color (scored 0–3), dryness (scored 0–4), and new foliage (scored 0–1), Krippendorff’s alpha was used to assess inter-rater reliability, with values of 0.8 or more indicating an acceptable level of agreement and α ≥ 0.667 serving as the lowest conceivable limit where tentative conclusions were still acceptable (Krippendorff, 2004). These calculations were made using R version 4.1.2 for Windows and the kripp.alpha function (irr: Various Coefficients of Interrater Reliability and Agreement. R package version 0.84.1).

## Results

### Efficacy of Oxathiapiprolin as a Protective Treatment Applied to Kauri Seedlings via Soil Drench

A plant trial was conducted to assess the efficacy of oxathiapiprolin as a protective treatment when applied to kauri seedlings in a soil drench before inoculation with the pathogen. Kauri seedlings were treated protectively with 10 or 50 mg oxathiapiprolin by diluting Zorvec® Enicade® with water and pouring it in the root zone of the seedling seven days prior to inoculation with *P. agathidicida* NZFS 3770. The seedlings were re-inoculated the following two weeks for a total of three inoculations. Six weeks after the first inoculation, five blinded, independent assessors rated wilting, colour, dryness, and new foliage in the seedlings (**Figure 1; Supplementary Materials Figure S1**), and these criteria were used to infer seedling health: scores were assumed to increase with disease severity. Krippendorff’s alpha was used to assess inter-rater reliability, with values of 0.8 or more indicating an acceptable level of agreement and α ≥ 0.667 serving as the lowest conceivable limit where tentative conclusions were still acceptable (Krippendorff, 2004). Agreement (ordinal scale assumed) was reached for wilting (α = 0.748), colour (α = 0.891), and dryness (α = 0.806). Therefore, the average score of the five assessors was used for analysis. New foliage had Krippendorff’s α = 0.580 (nominal scale assumed). However, these results were still reported, as only one rater appeared to differ most of the time, and one seedling was a particularly difficult determination: of the 15 cases, 5/5 raters agreed on 8 cases, 4/5 raters agreed on 6 cases, and the split was 2/3 for the difficult case.

**Figure 1.**
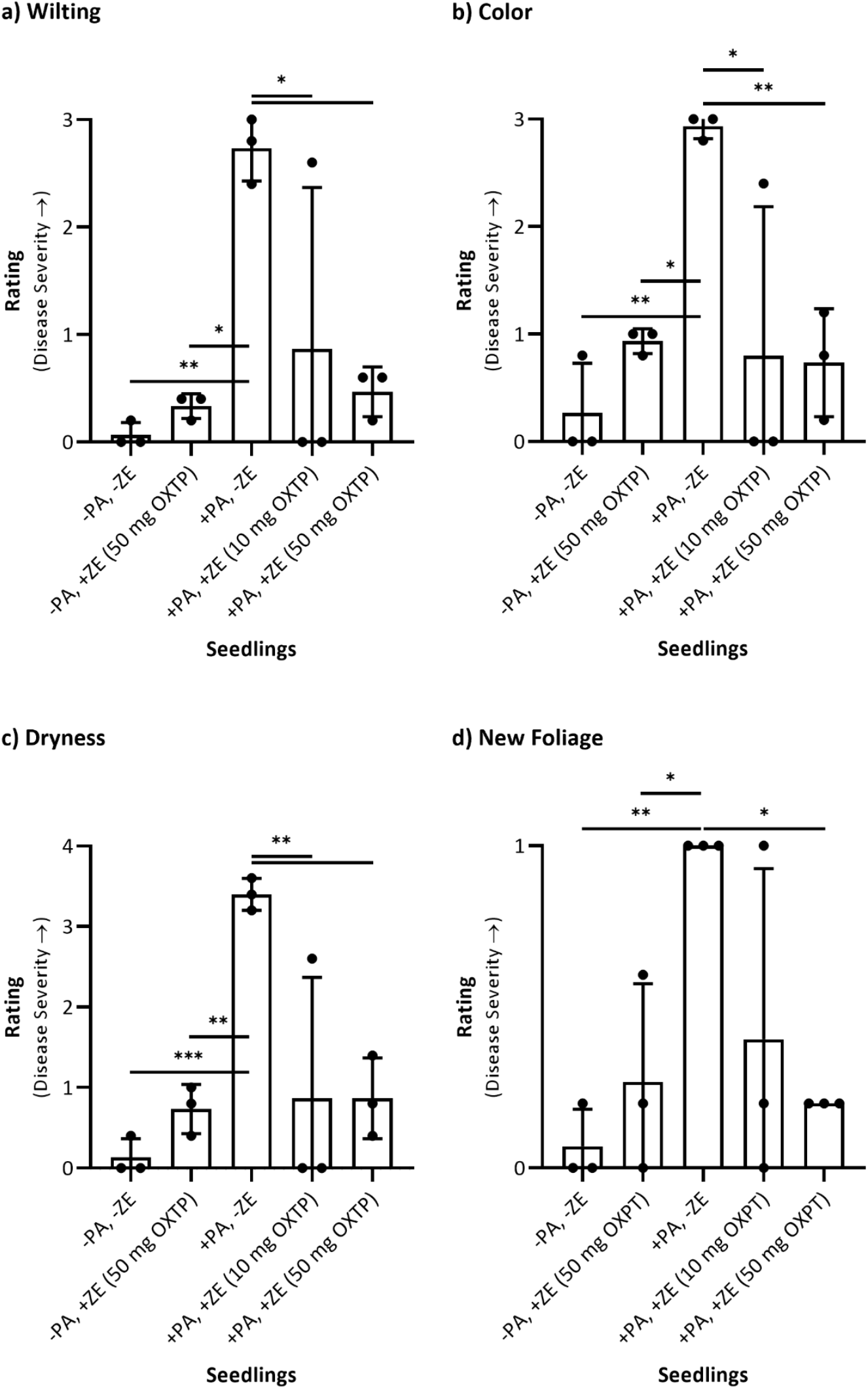
Results of the 5 blinded, independent ratings of a) wilting, b) color, c) dryness, and d) new foliage in the kauri seedlings treated protectively with Zorvec® Enicade® (10 or 50 mg oxathiapiprolin/seedling). The bars represent the mean (*n* = 3) ± standard deviation. To determine if the means from the data sets were significantly different from each other, a 2-way factorial ANOVA with the Tukey *post hoc* correction was conducted, and the difference was considered to be significant when the *p*-value was ≤ 0.05 (^*^*p* ≤ 0.05, ^**^*p* ≤ 0.01, ^***^*p* ≤ 0.001). Unless annotated, the difference was not considered to be significant. ± PA = inoculated or not inoculated with *P. agathidicida* NZFS 3770. ± ZE = treated or not treated with Zorvec® Enicade®. OXTP = oxathiapiprolin.

The ratings were significantly higher (*p* < 0.05) for the wilting, colour, dryness, and new foliage categories in the seedlings inoculated with *P. agathidicida* but untreated (+PA, -ZE) compared with the uninoculated seedlings (-PA), thus demonstrating the ability of *P. agathidicida* to cause dieback disease in untreated kauri seedlings. There were no significant differences in the ratings for all categories between the uninoculated/untreated seedlings (-PA, -ZE) and the uninoculated/treated seedlings (-PA, +ZE), meaning that the application of Zorvec® Enicade® (50 mg oxathiapiprolin/seedling) did not appear to affect seedling health (*i*.*e*. the treatment was not phytotoxic). The ratings were significantly lower (*p* < 0.05) for the wilting, colour, and dryness categories in the inoculated/treated seedlings (+PA, +ZE) compared to the inoculated/untreated seedlings (+PA, -ZE). For seedlings treated with 50 mg oxathiapiprolin (+PA, +ZE (50 mg OXTP)), there was also a significant increase (*p* < 0.05) in the presence and health of new foliage. Therefore, it was concluded that Zorvec® Enicade® (10 and 50 mg oxathiapiprolin/seedling) applied as a soil drench protected the kauri seedlings against infection by *P. agathidicida*, with the higher dosage being more effective.

### Efficacy of Oxathiapiprolin as a Curative Treatment Applied to Kauri Seedlings via Soil Drench

Similarly, a second plant trial was conducted to determine if oxathiapiprolin could be a curative treatment when applied to kauri seedlings in a soil drench after inoculation. On the day following the third and final inoculation with *P. agathidicida* NZFS 3770 (*i*.*e*. 15 days after the first inoculation), kauri seedlings were treated with dilute Zorvec® Enicade® to deliver 10 or 50 mg oxathiapiprolin *per* seedling. At that time, six of the nine infected kauri seedlings had possible wilting of new foliage, an early indication of disease. As before, five blinded, independent assessors rated wilting, colour, dryness, and new foliage six weeks after the first inoculation (**Figure 2; Supplementary Materials Figure S2**). Regarding inter-rater reliability, agreement (ordinal scale assumed) was reached for wilting (α = 0.798), colour (α = 0.704), dryness (α = 0.766), and new foliage (α = 0.886). Therefore, the average score of the five assessors was used for analysis.

**Figure 2.**
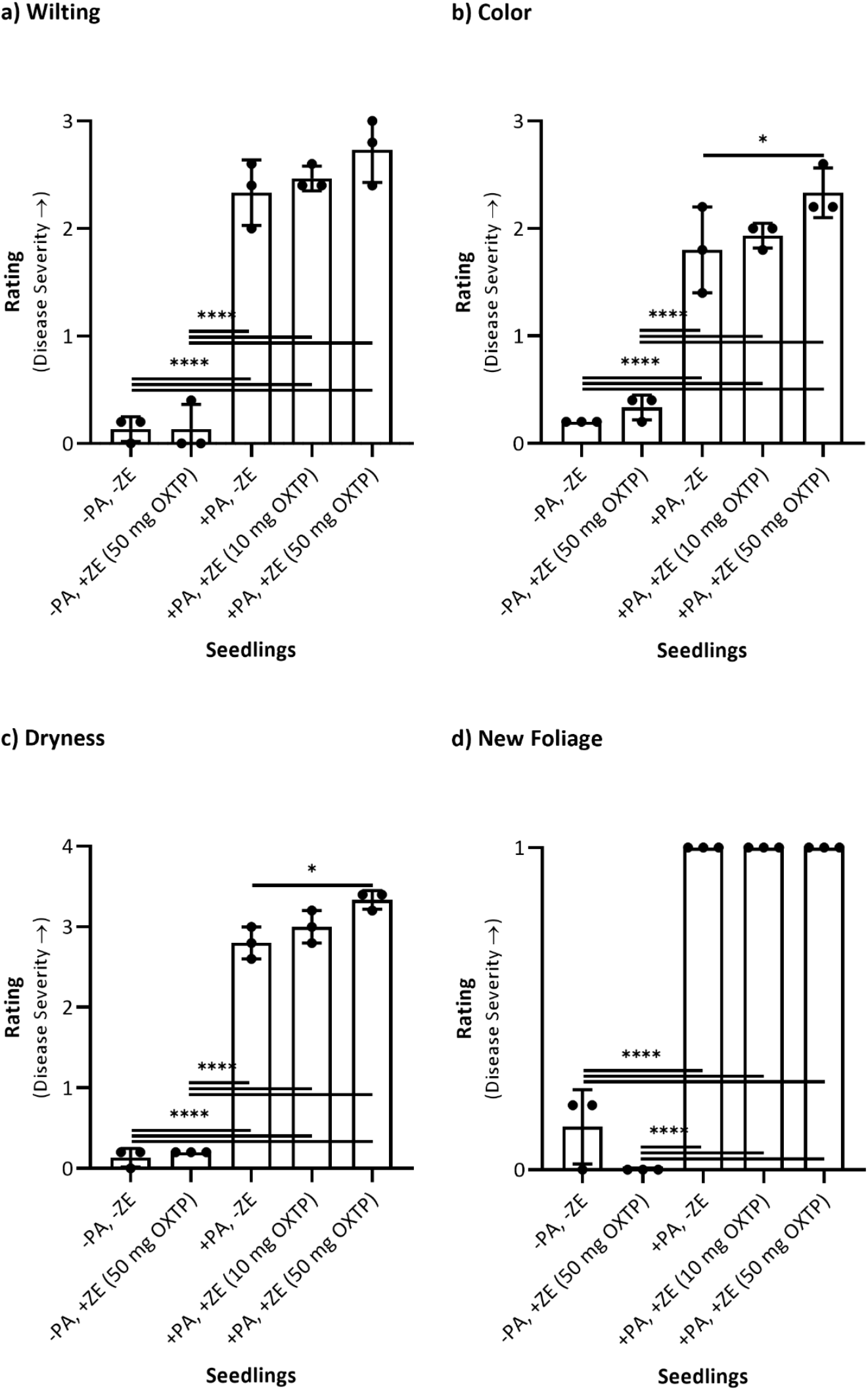
Results of the 5 blinded, independent ratings of a) wilting, b) color, c) dryness, and d) new foliage in the kauri seedlings treated curatively with Zorvec® Enicade® (10 or 50 mg oxathiapiprolin/seedling). The bars represent the mean (*n* = 3) ± standard deviation. To determine if the means from the data sets were significantly different from each other, a 2-way factorial ANOVA with the Tukey *post hoc* correction was conducted, and the difference was considered to be significant when the *p*-value was ≤ 0.05 (^*^*p* ≤ 0.05, ^**^*p* ≤ 0.01, ^***^*p* ≤ 0.001, ^****^*p* ≤ 0.0001). Unless annotated, the difference was not considered to be significant. ± PA = inoculated or not inoculated with *P. agathidicida* NZFS 3770. ± ZE = treated or not treated with Zorvec® Enicade®. OXTP = oxathiapiprolin.

Again, there were no significant differences in the ratings for all categories between the uninoculated/untreated seedlings (-PA, -ZE) and the uninoculated/treated seedlings (-PA, +ZE), meaning that the treatment (Zorvec® Enicade®, 50 mg oxathiapiprolin/seedling) did not appear to be phytotoxic. This assessment is congruous with the results of these controls in the protective plant trial. The disease ratings for the wilting, colour, dryness, and new foliage categories were significantly higher (*p* < 0.05) for the seedlings inoculated with *P. agathidicida* (+PA), whether treated or untreated (±ZE), compared to the uninoculated seedlings. There were significant differences (*p* < 0.05) in colour and dryness between the inoculated/untreated seedlings (+PA, -ZE) and the inoculated seedlings treated with 50 mg oxathiapiprolin (+PA, +ZE (50 mg OXTP)), with the treated seedlings showing more severe disease. Therefore, it was concluded that Zorvec® Enicade® (10 and 50 mg oxathiapiprolin/seedling) applied as a soil drench was not a curative treatment for the kauri seedlings already infected with *P. agathidicida*.

## Discussion

The experimental design of these plant trials (*e*.*g*. inoculation and treatment application methods, treatment amount) was based on the results of previous method development trials, as well as plant trials with oxathiapiprolin or kauri in the literature (Horner and Hough, 2013; Hao *et al*., 2019; Byers *et al*., 2021; Davison and Hardy, 2023). As in Byers *et al*. (2021), we inoculated kauri seedlings once a week for three weeks, thus ensuring sustained infection, by pouring mycelial mats into holes in the soil within the seedling root zone and saturating the soil to promote zoospore motility. The initial indicator of disease was the wilting of new foliage. Disease progression was characterised by leaves becoming increasingly wilted and dry and changing colour from green to yellow (*i*.*e*. chlorosis), brown, and orange. By six weeks following the first inoculation, all seedlings with dieback disease were dead or severely symptomatic, demonstrating the virulence of *P. agathidicida*. Consequently, this time point was selected as the end of the plant trials.

Similarly, six weeks after infection by *P. agathidicida* NZFS 3770, Byers *et al*. (2021) observed dieback symptoms in 18-month-old kauri seedlings, including chlorosis, wilting, leaf litter, necrosis, and mortality.

It was decided that the application of oxathiapiprolin as a soil drench was more likely to be effective than its application as a foliar spray because *P. agathidicida* is a root rot pathogen and oxathiapiprolin has systemic, acropetal movement in the xylem but no movement in the phloem (*i*.*e*. treatment moves up not down) (Pasteris *et al*., 2016, 2019). Horner and Hough (2013) also found that applying phosphite to kauri seedlings as a soil drench improved root and foliar health and tree survival for seedlings soil-inoculated with *P. agathidicida*. In contrast, foliar sprays were ineffective. Therefore, we applied oxathiapiprolin as a soil drench in these plant trials, but alternative application methods may be of interest in future studies.

In a method development experiment, we initially tested a range of amounts of oxathiapiprolin *per* kauri seedling (1, 10, 50, 100, 250, 500 mg) applied as a soil drench to determine the optimal dosage and found that 10 mg was the lowest amount to demonstrate a protective effect against infection by *P. agathidicida* NZFS 3770 (**Supplementary Materials, Figure S3**). Therefore, we selected this amount, as well as the next highest amount of 50 mg, for plant trials moving forward. Although oxathiapiprolin has potent *in vitro* activity against multiple lifecycle stages of *P. agathidicida* at sub-microgram *per* millilitre concentrations (Lacey *et al*., 2021), the necessity to upscale and apply milligram quantities of oxathiapiprolin to the seedlings was in agreement with expectations based on the literature: in these studies, microgram-to-milligram quantities of oxathiapiprolin were applied to field or vegetable crop plants, such as tomato, tobacco, and pepper plants, to prevent and/or cure infection by *Phytophthora* species (Ji *et al*., 2014; Matheron and Porchas, 2015; Miao *et al*., 2016; Bittner and Mila, 2017; Cohen, Rubin and Galperin, 2018; Cohen, 2020; Cohen and Rubin, 2020). More closely aligned with our work, Hao *et al*. (2019) performed greenhouse studies of 6–9 month-old citrus seedlings (‘Madam Vinous’ sweet orange) inoculated with root rot pathogens *Phytophthora nicotianae* or *Phytophthora citrophthora* and treated one week later with a soil drench of 8 mg oxathiapiprolin/seedling, and they found that the incidence of root rot and the pathogen populations in the soil were reduced to zero or near zero.

Our work demonstrates that 10 mg or, more effectively, 50 mg oxathiapiprolin *per* seedling applied as a soil drench is protective against dieback disease in kauri seedlings soil-inoculated with *P. agathidicida* NZFS 3770, but it is not curative. Both protective and curative effects of oxathiapiprolin against oomycete infection in plants have been reported (Ji *et al*., 2014; Matheron and Porchas, 2014, 2015; Ji and Csinos, 2015; Miao *et al*., 2016; Bittner and Mila, 2017; Cohen, Rubin and Galperin, 2018, 2019; Salas *et al*., 2019; Hao *et al*., 2019; Cohen, 2020; Cohen and Rubin, 2020). However, the extent of these effects can vary. For instance, Miao *et al*. (2016) applied oxathiapiprolin to pepper plants 24 h pre-inoculation (protective) or post-inoculation (curative) with *Phytophthora capsici*. These treatments demonstrated increased control of *P. capsici* infection whether applied before or after inoculation, but the protective activity was reported to be greater than the curative activity. Cohen *et al*. (2018) also reported that curative application of oxathiapiprolin to tomato plants was much less effective than protective application in controlling infection by *Phytophthora infestans*. Furthermore, Zorvec™ is only recommended for use in a preventative manner (Luijks, 2017). In our curative plant trial, two-thirds of infected seedlings were already possibly symptomatic at the time of treatment. Therefore, it is probable that the infection was already too advanced to be easily cured, since disease symptoms in aboveground parts of the plant often appear long after significant damage has already been done to the roots and the pathogen has developed a high biomass (Ivic, 2010).

These results demonstrate that oxathiapiprolin is a promising treatment to protect kauri from dieback disease, but it is likely to be less efficacious as a curative treatment, depending upon the timing of application post-infection.

## Conclusions

Our results show that the application of oxathiapiprolin (10 or 50 mg/seedling) as a soil drench to kauri seedlings is protective against infection by *P. agathidicida* NZFS 3770 but not curative once dieback symptoms are present. Oxathiapiprolin could have limited usefulness as a protective treatment in plant nurseries and for private landowners, but its use in a forest setting is less practical, facing the same challenges as other chemical controls: the determination of the appropriate dosage and required frequency of application, avoiding phytotoxicity and off-target effects, and the feasibility of treatment application (*e*.*g*. amount needed to cover large trees, cost, method, reaching the forest sites for repeated treatments). Before testing the capability of oxathiapiprolin to protect larger trees, however, additional seedling trials will need to be completed to assess different application methods and *P. agathidicida* isolates, as well as the protective activity on a longer timescale with multiple re-inoculations. Oxathiapiprolin should also be tested as a curative treatment when applied before the first disease symptoms. Furthermore, since oxathiapiprolin is at high risk for the development of resistance due to its single-site mode of action, it would be prudent to evaluate the activities of next-generation dual treatments that combine oxathiapiprolin with other oomycides (*e*.*g*. mefenoxam, famoxadone, benthiavalicarb, mandipropamid) to combat resistance.

## Author Contributions

**Hannah Robinson:** Methodology (lead), Formal analysis (lead), Investigation (lead), Supervision (supporting), Validation (lead), Visualization (lead), Writing – original draft (lead), Writing – review & editing (lead); **Jade Palmer:** Investigation (supporting), Writing – review & editing (supporting); **Monica Gerth:** Conceptualization (lead), Methodology (supporting), Funding acquisition (lead), Project administration (lead), Resources (lead), Supervision (lead), Writing – review & editing (supporting).

## Acknowledgements

The authors thank Lisa Woods for her support in the statistical analyses of the data, including her calculations of the inter-rater reliability. The authors also thank Corteva™ Agriscience (New Plymouth, NZ), for providing the Zorvec® Enicade® for our research. This research was supported *via* funding from Ngā Rākau Taketake (part of the BioHeritage National Science Challenge funded by the New Zealand Ministry of Business, Innovation and Employment (MBIE)) and MBIE Smart Ideas grant RTVU2201.

## Ethics Statement

None required.

