## Supplementary Materials for "Assessing the *in planta* efficacy of oxathiapiprolin as a potential treatment for kauri dieback disease"

### Supplementary Material

**Figure S1. Kauri seedlings on days 0 and 43 treated protectively with Zorvec® Enicade® (10 or 50 mg oxathiapiprolin/seedling).** Brightness, contrast, and saturation adjusted in the pictures for better visualization of effect. BR = biological replicate. ± PA = inoculated or not inoculated with *P. agathidicida* NZFS 3770. ± ZE = treated or not treated with Zorvec® Enicade®. OXTP = oxathiapiprolin.

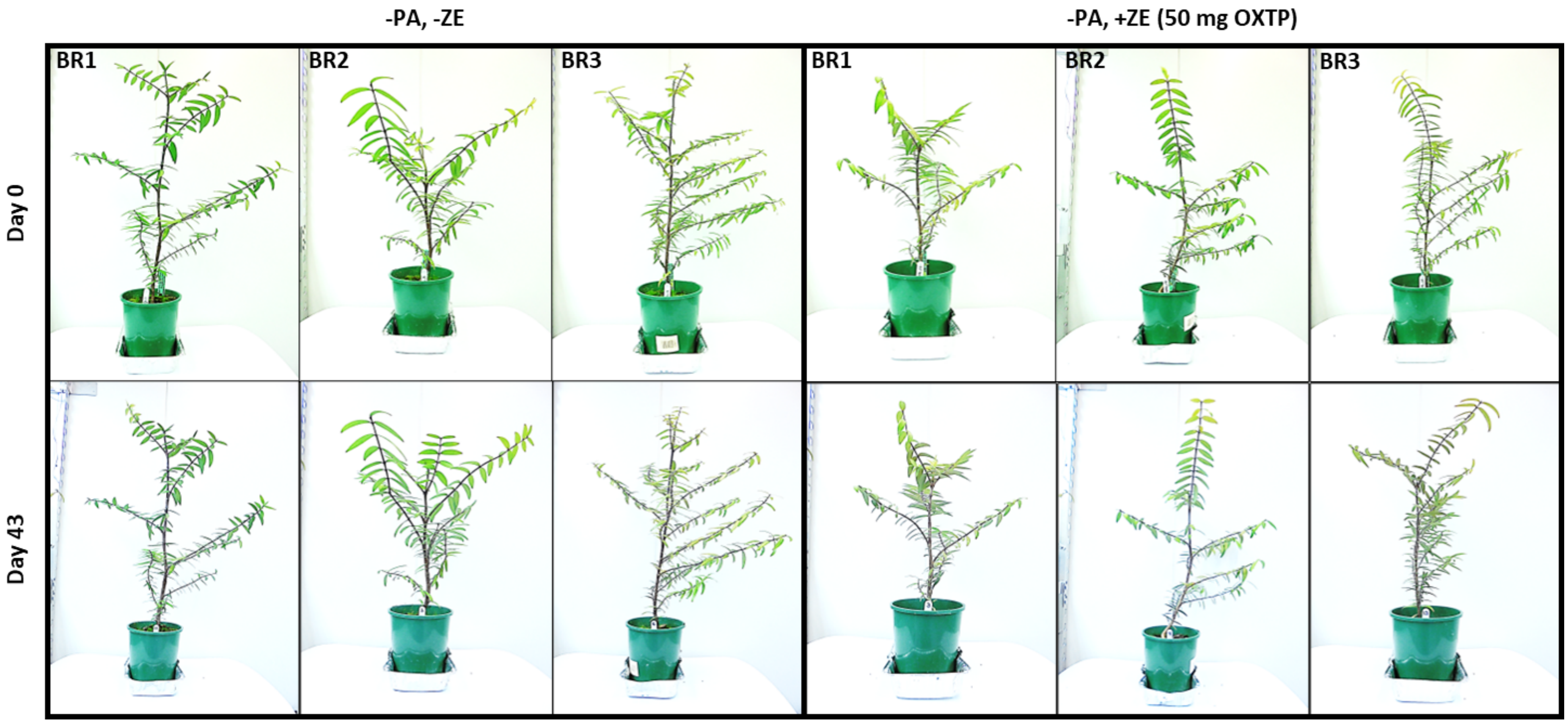

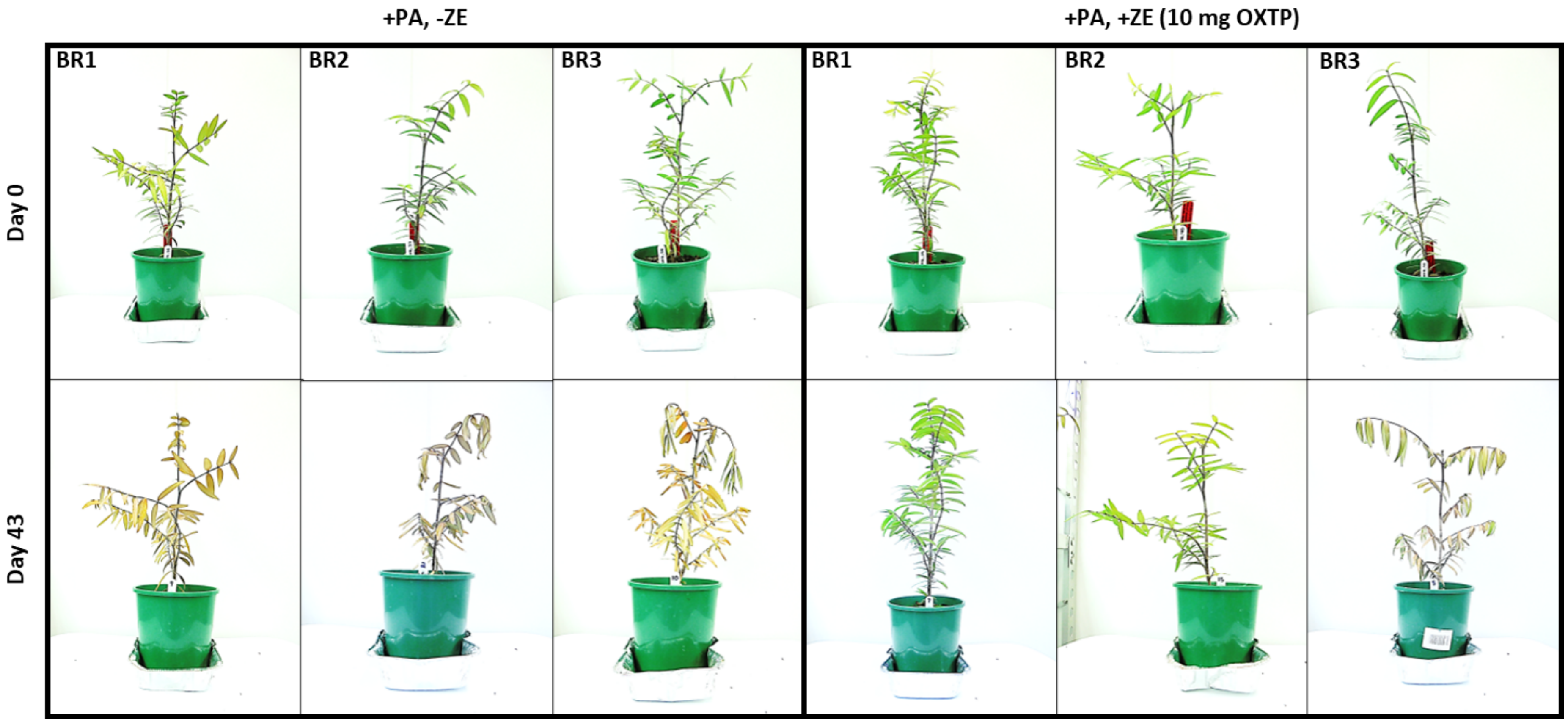
+PA, -ZE
+PA, +ZE (10 mg OXTP)
BR1
BR2
BR3
BR1
BR2
BR3
Day 0
Day 43

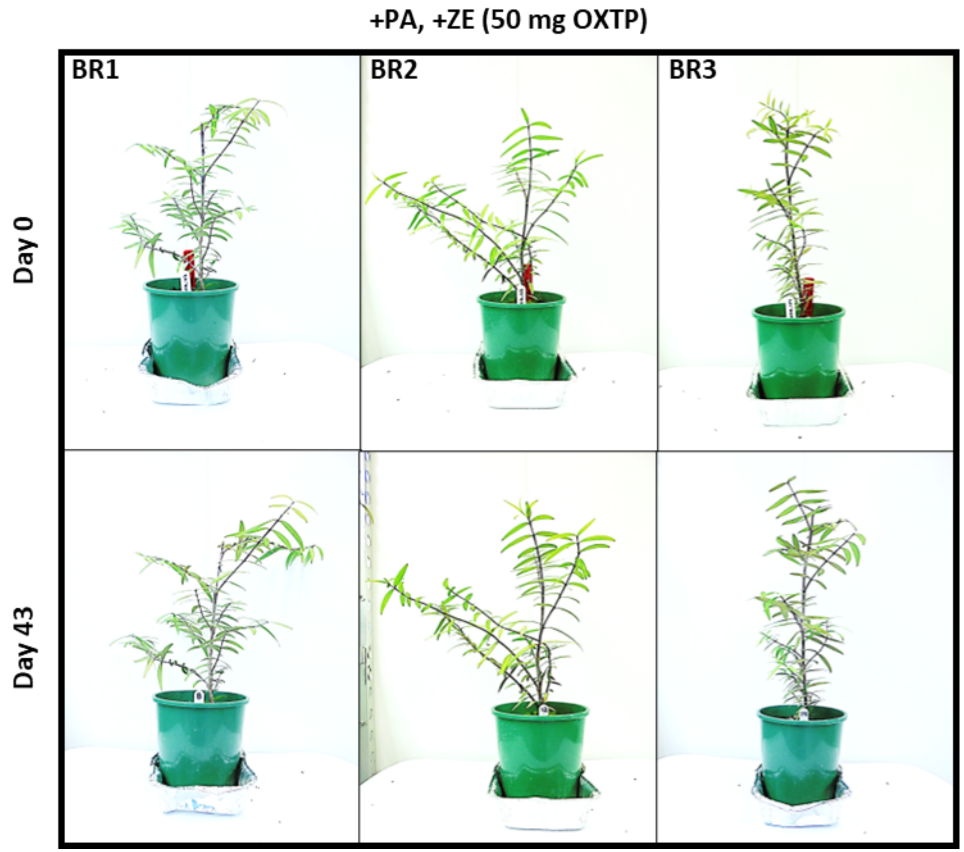
+PA, +ZE (50 mg OXTP)
BR1
BR2
BR3
Day 0
Day 43

**Figure S2. Kauri seedlings on days 0 and 42 treated curatively with Zorvec® Enicade® (10 or 50 mg oxathiapiprolin/seedling).** Brightness, contrast, and saturation adjusted in the pictures for better visualization of effect. BR = biological replicate. ± PA = inoculated or not inoculated with *P. agathidicida* NZFS 3770. ± ZE = treated or not treated with Zorvec® Enicade®. OXTP = oxathiapiprolin.

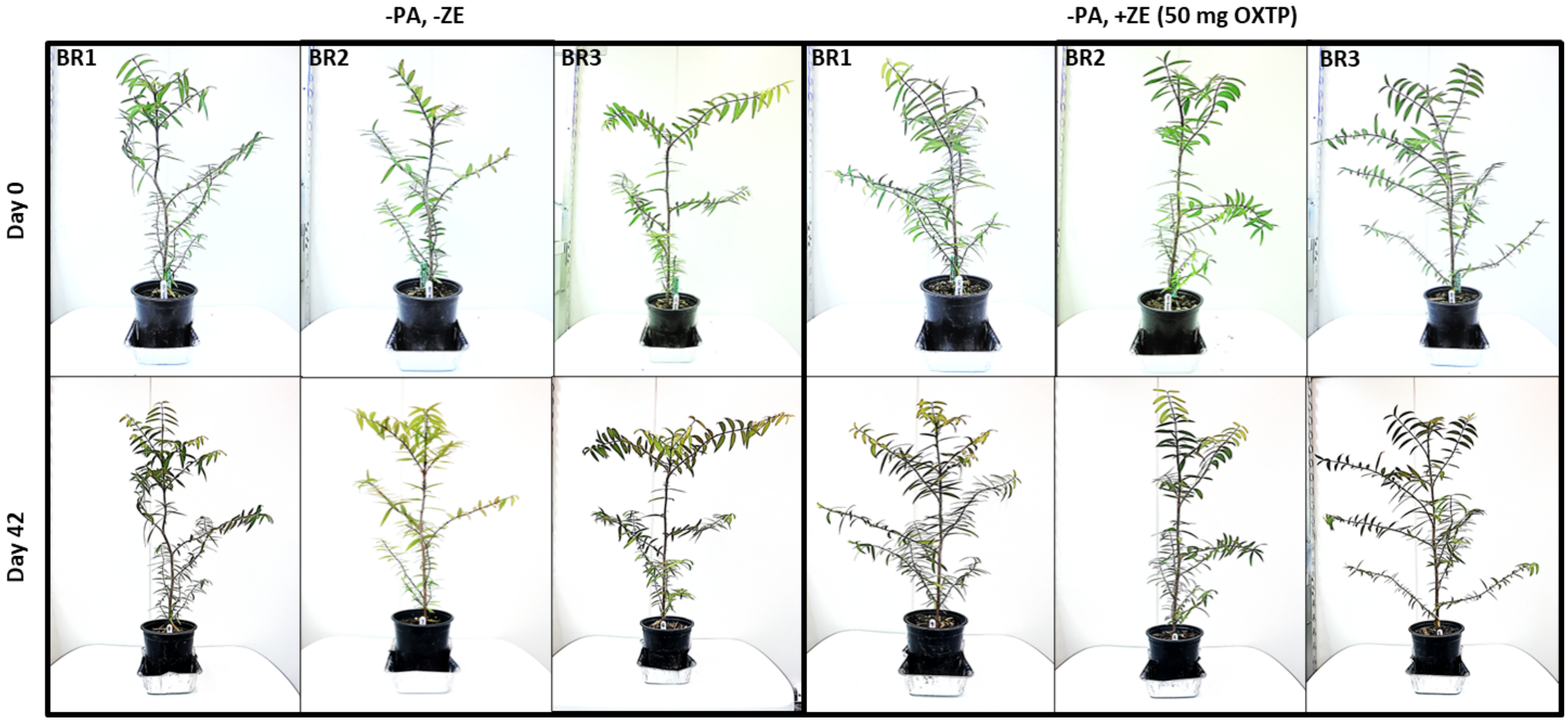

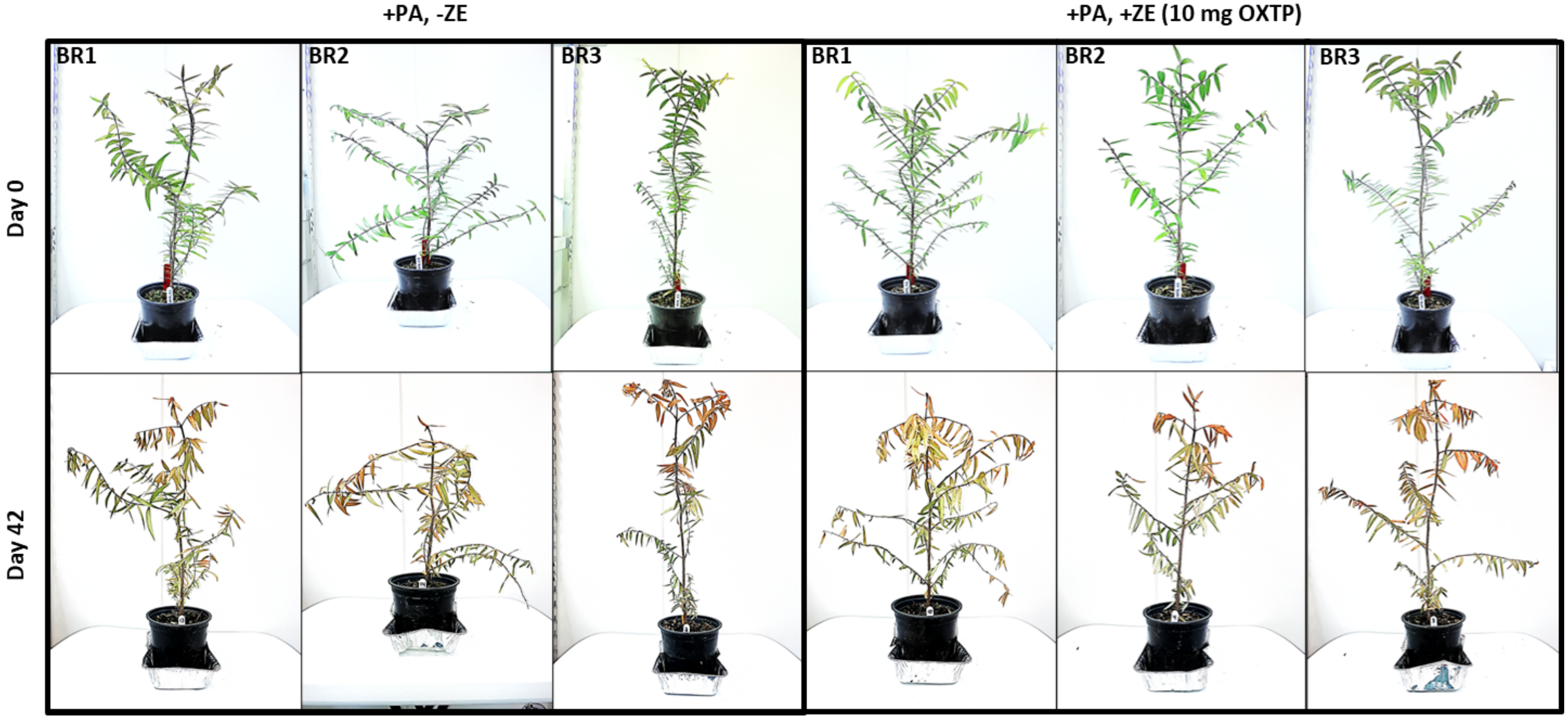
+PA, -ZE
+PA, +ZE (10 mg OXTP)
BR1
BR2
BR3
BR1
BR2
BR3
Day 0
Day 42

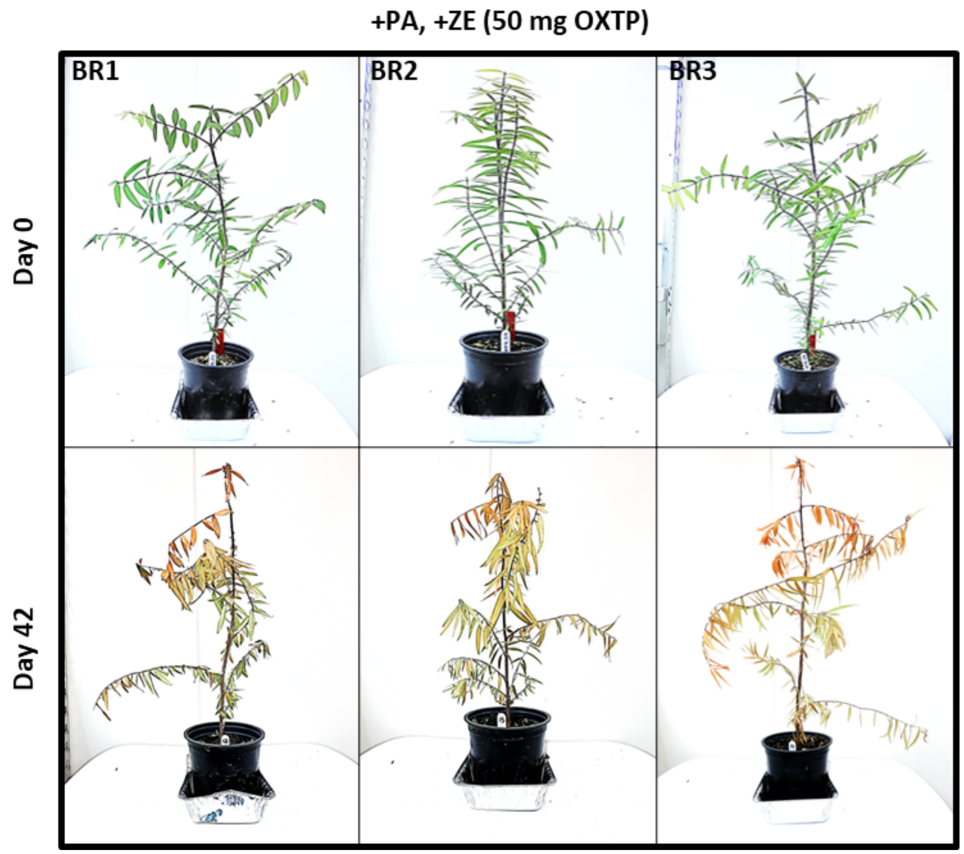
+PA, +ZE (50 mg OXTP)
BR1
BR2
BR3
Day 0
Day 42

**Figure S3. Kauri seedlings on days 0, 44, and 109 treated protectively with Zorvec® Enicade® (1, 10, 50, 100, 250, or 500 mg oxathiapiprolin/seedling).** Brightness, contrast, and saturation adjusted in the pictures for better visualization of effect. ± PA = inoculated or not inoculated with *P. agathidicida* NZFS 3770. ± ZE = treated or not treated with Zorvec® Enicade®. OXTP = oxathiapiprolin.

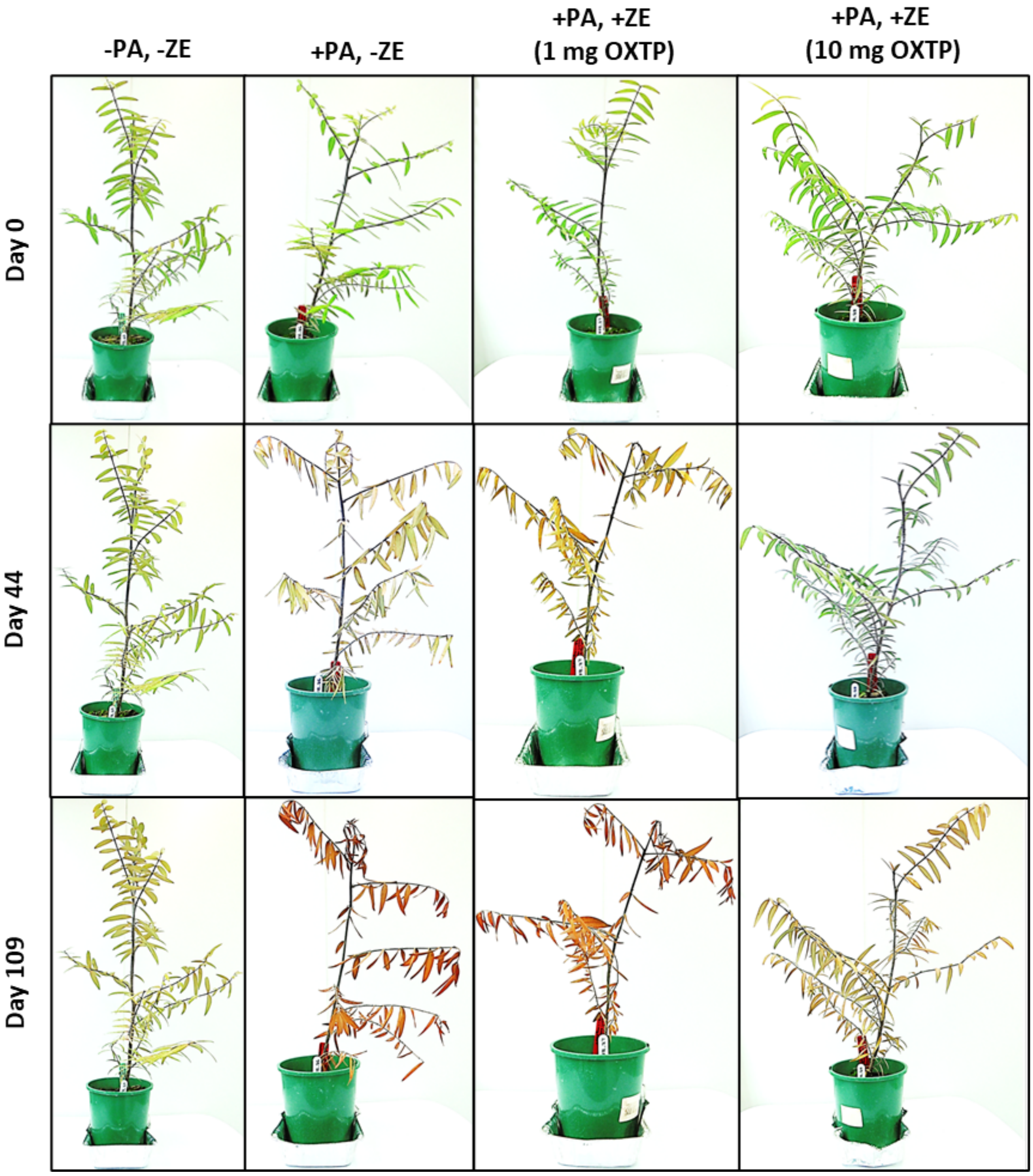

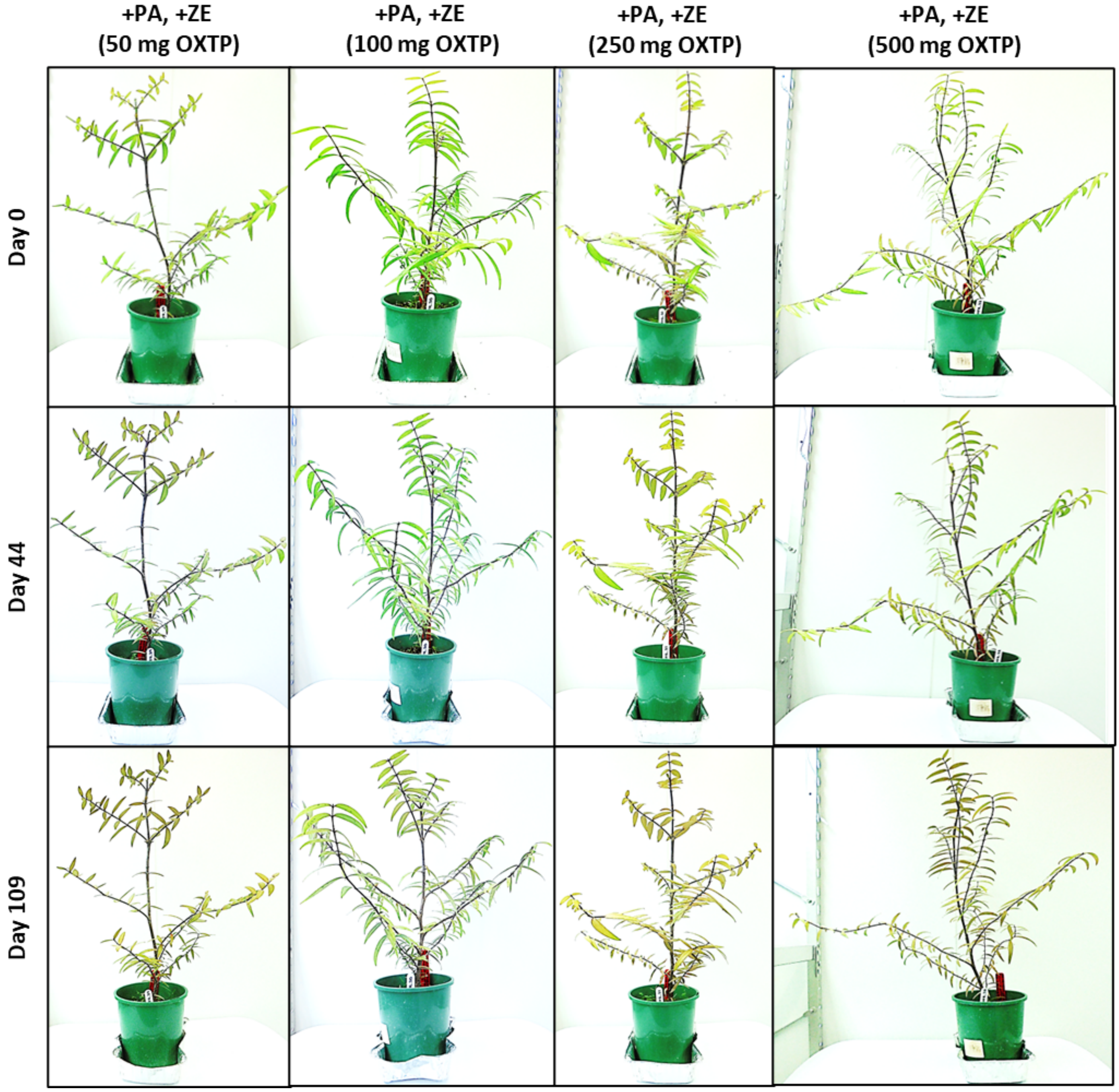
+PA, +ZE
(50 mg OXTP)
+PA, +ZE
(100 mg OXTP)
+PA, +ZE
(250 mg OXTP)
+PA, +ZE
(500 mg OXTP)
Day 0
Day 44
Day 109
